# Distinct molecular profile of the chick Organizer as a stem zone during axial elongation

**DOI:** 10.1101/2022.12.16.520706

**Authors:** Timothy R. Wood, Iwo Kucinski, Octavian Voiculescu

**Affiliations:** Department of Physiology, Development and Neuroscience, University of Cambridge, Downing Street, Cambridge CB2 3DY, UK; Department of Haematology, Cambridge Institute for Medical Research, University of Cambridge, Hills Road, Cambridge CB2 0XY, UK

**Keywords:** stem zone, organizer, axial elongation, WIF1, PTGDS, ThPO, UCKL1, chick

## Abstract

The vertebrate Organizer plays a crucial role in defining the main axes of embryo: it neuralizes the surrounding ectoderm, and is the site of emigration for cells making axial and paraxial mesendoderm during elongation. The chick Organizer becomes a stem zone at the onset of elongation: it stops recruiting cells from the neighbouring ectoderm, and generates all its derivatives from the small number of resident cells it contains at the end of gastrulation stages. Nothing is known about the molecular identity of this stem zone. Here, we specifically labelled long-term resident cells of the Organizer, and compared their RNA-seq profile to that of the neighbouring cell populations. Screening by RT-PCR and in situ hybridisation identified four genes (*WIF1, PTGDS, ThPO* and *UCKL1*), which are upregulated only in the Organizer region when it becomes a stem zone, and remain expressed there during axial elongation. In experiments specifically labelling the resident cells of the mature Organizer, we show that only these cells express these genes. These findings molecularly define the Organizer as a stem zone, and offer a key to understanding how this zone is set up, the molecular control of its cells’ behaviour, and the evolution of axial growth zones.

## Introduction

The anterior-posterior axis of all vertebrate embryos is formed by a combination of two mechanisms: direct specification of anterior structures, and sequential addition of material generated by posterior growth zones. Both processes involve a small region of the gastrulation site, the Organizer. In Amniotes, gastrulation takes place along the primitive streak (PS), with the Organizer (node) at its tip. In the chick embryo, it has been shown that the Organizer (Hensen’s node) acts in distinct ways during gastrulation and elongation stages.

During gastrulation, the Organizer (as the rest of of the primitive streak) constantly changes its cellular composition. The epithelial cells at the PS undergo epithelial-to-mesenchymal transition (EMT) and ingress into lower layers; the constriction of their apical surfaces causes a pull on the cells in the neighbouring epiblast into the PS, which are in turn induced to undergo EMT^1^. At gastrulation stages, the Organizer mainly produces gut endoderm^2,3^. A sharp transition marks the end of gastrulation and the beginning of axis elongation (Figure 1, Supplementary Video 1): the Organizer stops recruiting cells^4^ as it starts laying down its derivatives^5^ in head-to-tail sequence (head mesoderm, notochord and somitic mesoderm). Also at the beginning of axial elongation, the ectoderm close to the Organizer is stably neuralized. The neural territory anterior to the Organizer is specified as brain, while the regions on the sides of the Organizer become growth zones that sequentially generate the caudal hindbrain and the spinal cord^6,7^.

**Figure 1:**
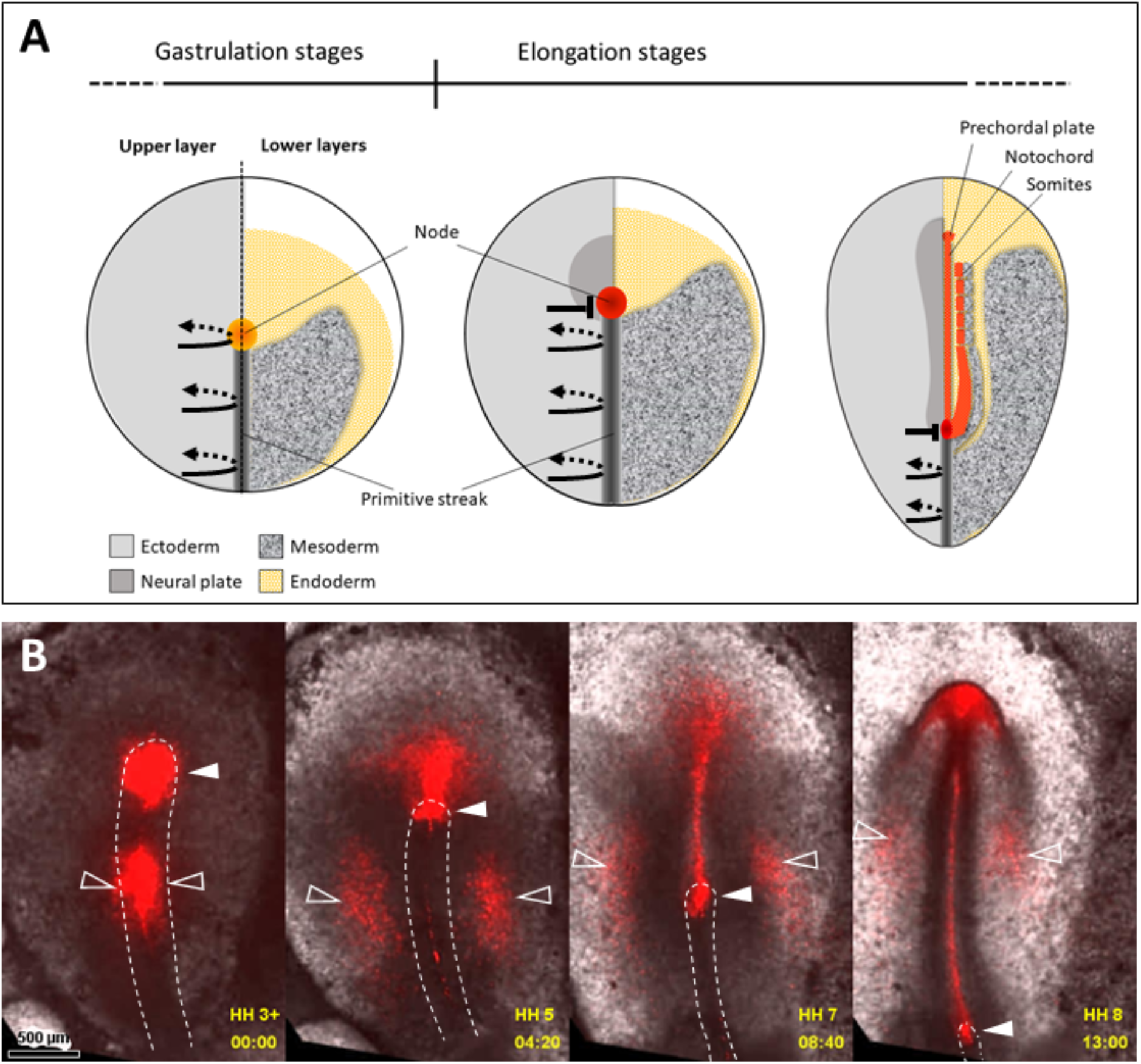
The chick Organizer acts as a stem zone generating axial and paraxial mesoderm during elongation. **(A)** Diagram summarising the existing fate maps. Embryos at successive stages are shown in a split view, with the upper layer on the left side and the lower layers on the right side. Recruitment of epiblast cells to the primitive streak (black curved arrows) ceases at the level of the Organizer at the initiation of elongation stages (black block sign). **(B)** Embryo with the Organizer and mid-primitive streak fluorescently labelled (open arrowheads), showing that the Organizer retains resident cells that generate axial and paraxial structures (solid arrowheads), while all cells in the rest of the primitive streak emigrate away from it. Frames from Supplementary Video 1. Anterior to the top, dorsal views.

Remarkably, the Organizer generates all its derivatives from the few cells it contains at the initiation of axial elongation. Single-cell labelling experiments have shown the Organizer contains long-term residents capable of self-renewal, which also generate descendants contributing to head mesendoderm, and axial and paraxial mesoderm^8,9^.

The molecular mechanisms setting up the Organizer at gastrulation are well understood and conserved across vertebrates^10,11^. In contrast, nothing is known about the molecular and cellular mechanisms controlling the maintenance and function of the Organizer during axial elongation. At the start of axial elongation, the Organizer stops expressing genes associated with gastrulation stages, and also loses its morphological definition, its edges becoming indistinguishable from surrounding neural ectoderm.

We therefore set out to identify genes specifically associated with the chick Organizer at axial elongation stages. To this end, we specifically labelled the Organizer stem zone and compared its transcriptome profile with that of adjoining regions. Next, we performed expression screenings by RT-PCR and in situ hybridisation for transcripts of higher expression in the mature Organizer. This led to the identification of four genes (*WIF1, PTGDS, ThPO* and *UCKL1*), which are upregulated in the Organizer at the start of elongation stages and remain expressed in this region as the axis is laid down. Graft experiments show that expression of these genes is specific to resident cells of the mature Organizer. Most remarkably, these genes code for molecules of unrelated classes that are not known to be co-expressed in any other tissue. This set of genes now offer an unambiguous definition of the mature Organizer as a stem zone, and can be used to distinguish it from adjacent stem cell populations. We discuss their possible functional significance, and their use in evo-devo analysis of stem populations fuelling axial elongation.

## Results

### A unique molecular profile of the mesodermal stem zone

We sought to identify genes specifically expressed by the Organizer as it becomes a mesodermal stem zone during elongation, and distinguish it from the accompanying stem zones fuelling the extension of the neural plate. To this end, we devised a method of labelling resident Organizer cells during axial elongation (Figure 2A, B). We fluorescently labelled the Organizer cells with a lipophilic dye (CM-DiI), just before it becomes a stem zone and the embryo starts elongating (fully extended PS, late stage HH 3^+^; see Figure 2A). Labelled embryos were then allowed to develop for 16-24 hours (Figure 2B). In order to obtain transcription profiles of node resident cells and select the transcripts whose expression does not change over time, cells were collected from embryos at a range of developmental stages, between 8-10 somites. Two separated cell populations were collected: (i) fluorescently labelled cells remaining in the anterior primitive streak (‘R’ in Figure 2 C, D) and (ii) adjacent non-labelled cells in the neural plate or paraxial mesoderm (‘W’ in Figure 2 C, D). We used two methods to achieve this: laser capture on transverse sections through the node (Figure 2 C) and manual dissection of fluorescent and non-fluorescent cells from the region of the anterior streak (Figure 2 D). We found each technique presented complementary advantages and drawbacks (see Methods): while manual dissection produces best RNA quality (RIN = 10) but is relatively coarse, laser capture offers more precise separation between the populations but the quality of RNA is lower (RIN values around 7). For each method, three pairs of biological replicates were deep sequenced and compared (Figure 2 E and Supplementary Table 1). We selected 93 transcripts that consistently showed higher gene counts in anterior primitive streak resident cells than in neighbouring cells (Supplementary Table 2). To examine their expression in the embryo, we first used RT-PCR on cDNA extracted from six different locations that included all embryonic layers (Figure 2 F): the anterior primitive streak, anterior midline containing some of the Organizer derivatives, two neural domains (caudal and anterior), the caudal primitive streak, and the non-neural ectoderm. This initial screen indicated that 32 transcripts are present only in cDNA from the anterior region of the primitive streak and the anterior midline (Supplementary Table 2). These transcripts were further examined by in situ hybridization. Markers of mature Organizer stem cells should fulfil the following criteria: (i) are not expressed during gastrulation, (ii) are only upregulated in the anterior primitive streak at the initiation of axial elongation, and (iii) their expression is retained at this location throughout elongation. We found four genes that meet these criteria: *cWIF1, cTPO, cPTGDS* and *cUCKL1* (Figure 3). The expression patterns of these genes in the anterior primitive streak correlates with the location of the mature Organizer resident cells. However, at the onset of their expression (end of gastrulation stages), the anterior primitive streak contains several resident populations as well as transient cells, which occupy overlapping domains. At subsequent stages, the Organizer also loses its morphological boundaries and it is difficult to distinguish it from the neighbouring growth zones of the neural plate. We therefore set out to test more accurately how specific are the expression patterns of the newly identified genes for the long-term resident cells of the Organizer.

**Figure 2:**
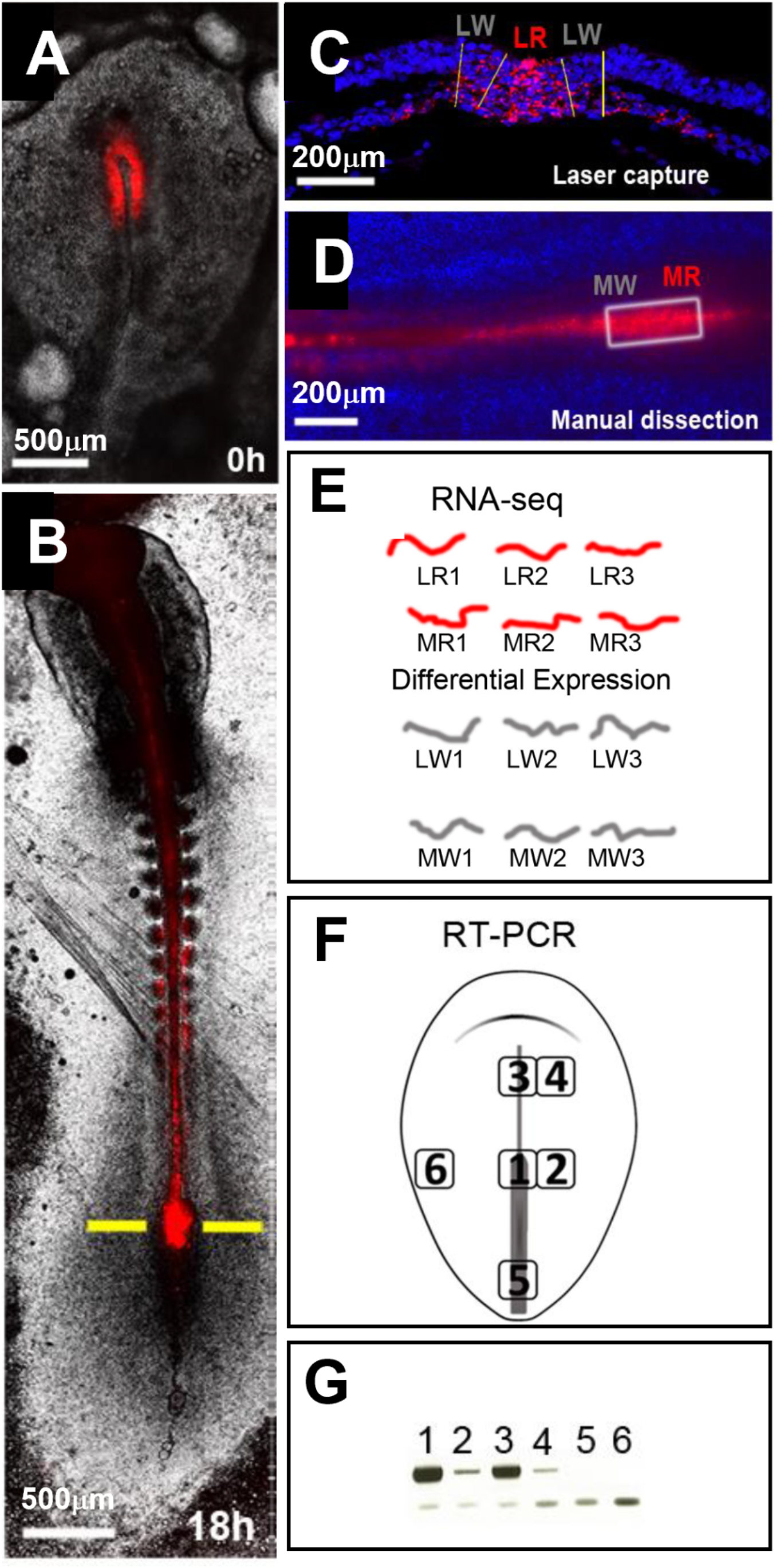
Method for the molecular profiling of Organizer resident cells. **(A, B)** Fluorescent labelling (red) the Organizer at the end of gastrulation (A) allows the selective marking of its resident cells during elongation stages (B). **(C)** Thin section at the level of the anterior primitive streak (yellow line in (B)), showing the labelled and unlabelled collected by laser microdissection. **(D)** Region containing the anterior primitive streak and surrounding regions, used for the manual dissection of labelled and unlabelled cells. **(E)** Diagram of RNA-seq samples used for differential gene counts analysis. **(F)** Embryonic regions used for RT-PCR screening. **(G)** Example of RT-PCR results showing specific expression in the anterior primitive streak and its derivatives. Lower bands are primers, upper bands are PCR products. LR: laser dissection resident cells; LW: laser dissection non-labelled cells; MR: manual dissection resident cells; MW: manual dissection non-labelled cells.

**Figure 3.**
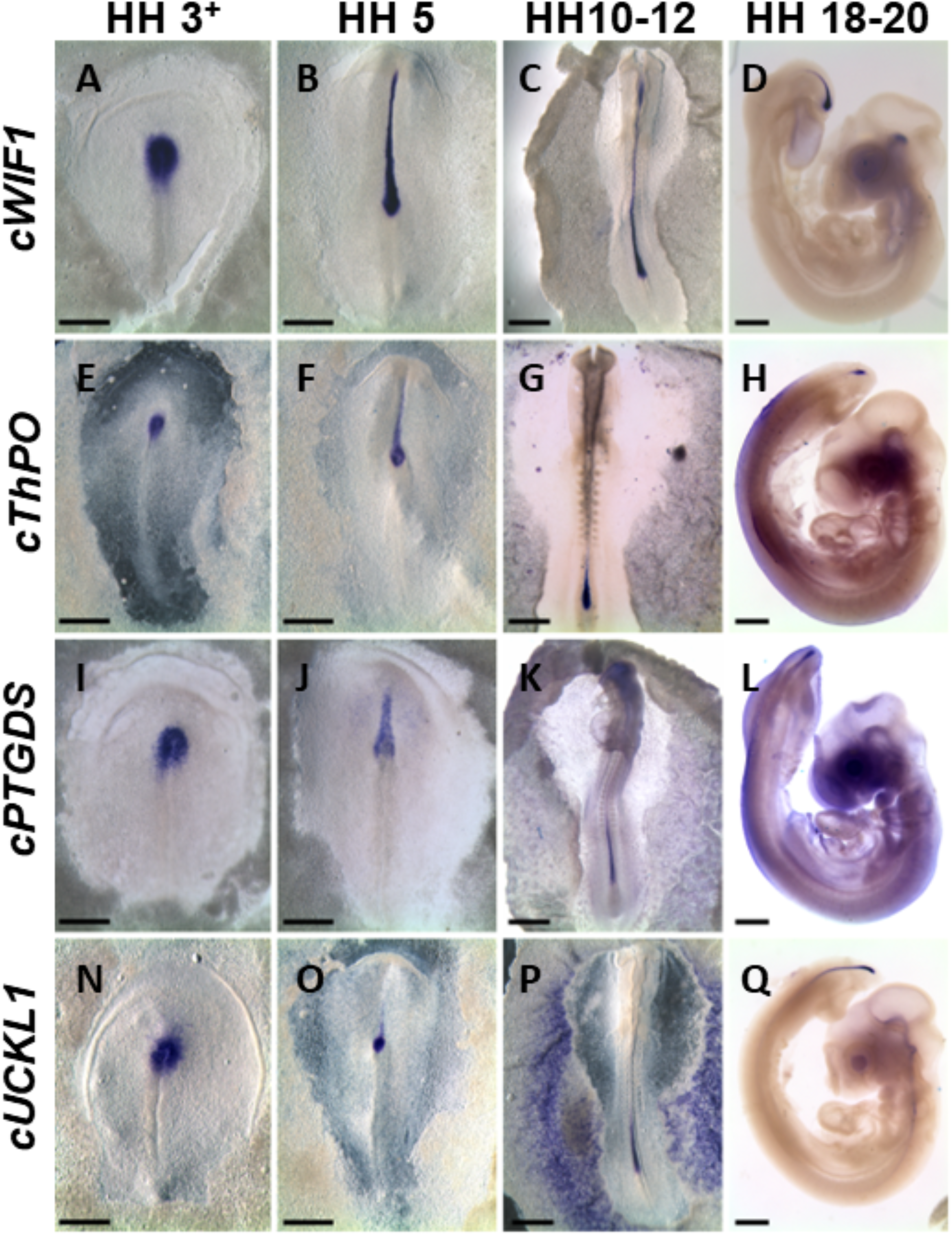
Genes expressed in the anterior primitive streak throughout elongation. In situ hybridisation for *Wif1* (Wnt Inhibitory Factor 1), *ThPO* (Thrombopoietin), *PTGDS* (Prostaglandin D2 Synthase), *UCKL1* (Uridine-Cytidine Kinase Like 1). Scale bars, 500 µm (A-C, E-G, I-K, M-P), 1 mm (D, H, L, Q).

### Resident node cells retain expression of mesodermal stem cells

To test whether these genes are specific for the long-term resident cells of the Organiser, all the cells in the anterior streak (both transient and long-term residents) need to be labelled at the initiation of elongation. Transient cells are then allowed to emigrate during subsequent stages, thus leaving the resident cells as the only labelled population. Marking all the cells of the Organizer was achieved by homotopic and isochronic grafts of the anterior primitive streak region, from fluorescently labelled embryos into unlabelled hosts at the initiation of elongation. Embryos were then allowed to develop for 18-24h to stages 5-15 somites (Figure 4 A-C). Immunohistochemistry (Figure 4 D, E) was employed to evidence all the cells bearing the fluorescent marker, and the expression pattern of the newly identified genes was assessed by in situ hybridization. In sections at the level of the anterior primitive streak, we could only identify double positive or double negative cells (Fig. 4 F, G). These results show that the genes identified here are bona fide markers of the resident anterior primitive streak cell population.

**Figure 4.**
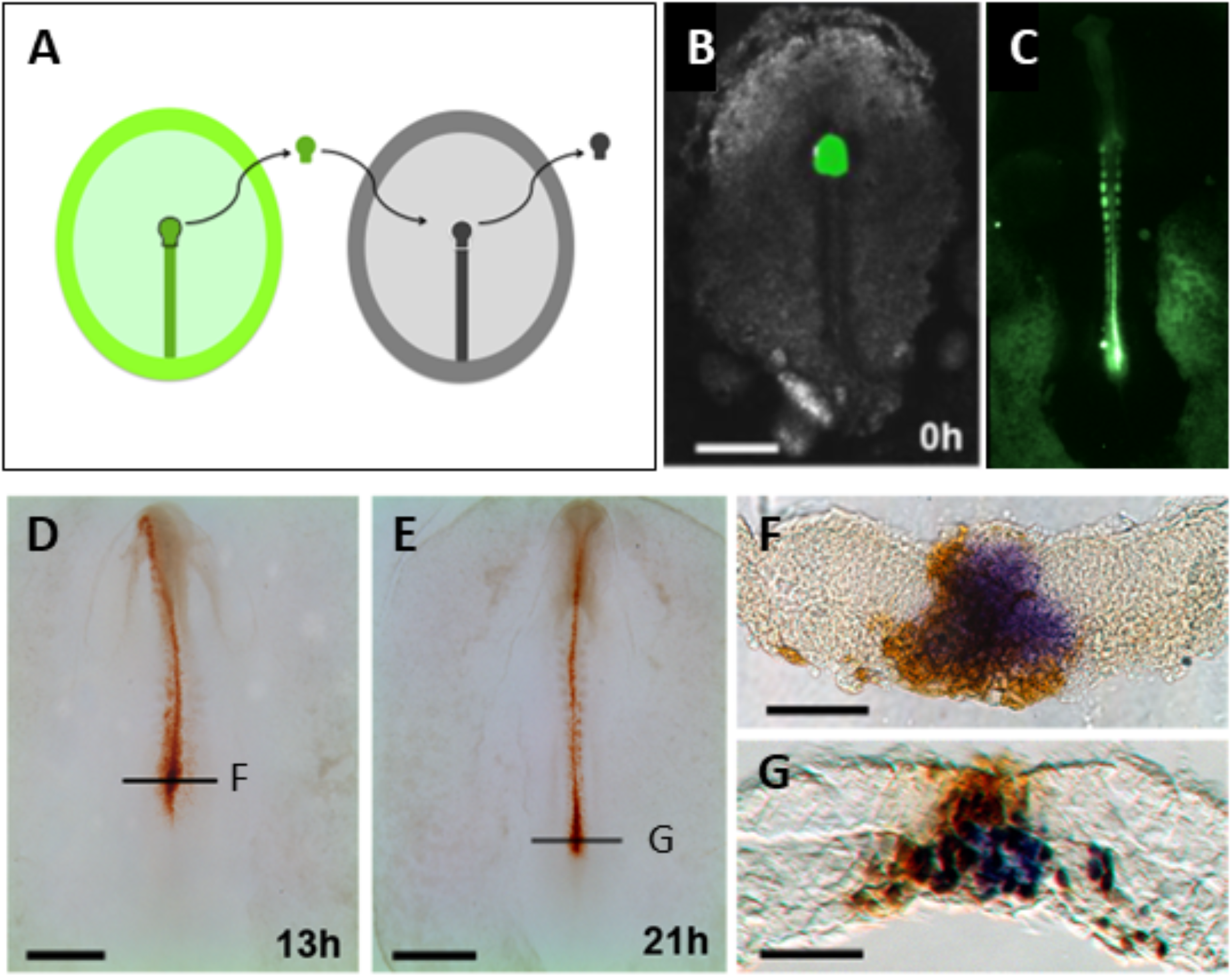
Specific expression of the identified genes in Organizer resident cells. **(A)** Homotopic and isochronic (HH3^+^) graft of a labelled anterior primitive streak to label all the resident cells in the Organizer. Example of grafted embryo with fluorescently labelled cells in green just after transplantation **(B)**, and after 24h of culture **(C). (D, E)** Examples of grafted embryos at different developmental stages, processed by immunohistochemistry for the fluorescent marker in brown. **(F, G)** Grafted embryos processed for both in situ hybridisation (*cWif1*, purple) and immunohistochemistry (brown). Thin sections at the levels indicated in (D, E).

## Discussion

Fate map experiments together with single-cell lineage analysis showed the presence of a mesodermal stem cell population at the mature Organizer during elongation^5–7,12^. However, the distinction of this stem cell population, as well as the study of its cellular behaviours and function, has been hampered by the lack of specific marker genes for this population. Here we identified a distinct molecular signature defining the chick Organizer as a stem zone during elongation stages. These findings now permit addressing several important issues.

First, this set of genes can be used to rigorously distinguish between resident Organizer cells (mesodermal stem zone) and adjacent neighbouring stem populations (caudal neural plate and nero-mesodermal bipotent progenitors).

Second, it is now possible to explore the molecular mechanisms regulating the mature Organizer cell behaviours. The four genes identified here are specific markers of the mesodermal stem zone (*WIF1, PTGDS, TPO* and *UCLK1*) and code for proteins of diverse classes and functions. Their known functions immediately suggest several hypotheses. WIF1 is an inhibitor of both canonical and non-canonical Wnt pathways, which are involved in regulating multipotentiality in other stem cell populations^13^. The prostaglandin synthase PTGDS has been implicated in controlling EMT^14,15^ in other cellular contexts. It has also been shown to trap and/or transport retinoids^16^, independently of its catalytic function. This suggests that PTGDS may be involved in protecting the Organizer cells from the action of retindoids produced in more anterior regions. Thrombopoietin controls proliferation in hematopoietic niches^17–19^, and UCKL1 in blast transformation^20,21^; thus, these genes may be involved in controlling cell cycle progression of mature Organizer progenitors.

Third, the regulation of these genes offers a key to investigate how the mature Organizer is established as a stem zone. To our knowledge, the four genes found here are not co-expressed in any other tissues and it seems unlikely that any of them directly regulates the expression of the others. Finding the gene regulatory network of these genes will shed light on the molecular mechanisms responsible for the acquisition of stem cell properties by the Organizer. Moreover, these markers can be used to understand what induces the stemness of the Organizer at the end of gastrulation stages. Is this induced by signals emanating from neighbouring structures at the beginning of elongation (e.g. more posterior primitive streak, neural plate), or does it result autonomously from the temporal progression of the Organizer?

Finally, how evolutionarily conserved is this signature of the mesodermal stem zone? In all vertebrate and most invertebrate embryos, the long axis is laid down through a combination of direct specification and posterior growth zones. It is still debated what are the relative contributions of these mechanisms, and how evolutionarily conserved growth zones are. Examining the expression of these genes’ homologues in other vertebrates will help to identify the mesodermal stem cells, how these cells are arranged with respect to other progenitors, and the relative contribution of growth zones to axial elongation across vertebrates.

## Materials and Methods

Fertilized eggs of wild-type hens (Bovans Brown) were obtained from WinterEgg Farm (Thriplow, Herts, UK).

### Embryo culture

Fresh, fertilised hen’s eggs were kept for up to one week at 15°C until use, and incubated at 38°C for the embryo to reach the desired stages. Embryos were explanted and manipulated in Pannett-Compton saline, and cultured using New’s technique^22^, as modified and described previously^23^.

### Fluorescent labelling

Working dilutions of lipophilic CM-DiI (Invitrogen, cat. # C7001) was prepared fresh, by adding 1µl of stock solutions (0.5% w/v in ethanol, kept at -20°C) to 9µl of aqueous sucrose solution (6% w/v) pre-warmed to 37°C. Finely drawn capillaries, with a tip broken to a bore of about 1-2 µm, were used to deposit warm solution close to the epiblast of embryos kept in saline.

### Microscopy

Time-lapse epifluorescence movies were acquired with a PlanApo N 2x/0.08 lens on an Olympus IX71 wide-field microscope, fitted with fluorescence excitation source (CoolLed pE-2) and appropriate emission optics, motorised stage (Prior), and Hamamatsu C8484 camera controlled by HCImage software. The system was enclosed in a Perspex box, one wall of which is the top of a Marsh Automatic Incubator (Lyon, U.S.A.) to provide thermoregulation and air circulation.

### Whole node transplantations

Donor embryos were fluorescently labelled by CMFDA (Invitrogen, cat. # C2925) by bathing whole, for 1 hour at 38ºC, in Pannett-Compton solution containing CMFDA diluted 1:250 from stock solution (10 mM in DMSO, kept at -20ºC). They were then transferred three times for 10 minutes each in fresh Pannett-Compton saline to remove excess dye, before having their nodes excised. The nodes were excised from unlabelled receiving embryos, prepared in New culture; the labelled nodes were aspirated and transferred with the aid of a P2 pipette adjusted to 0.2 µl.

### Microsurgery

Surgical manipulations were done using fine tungsten needles sharpened by electroelution ^57^. For RNA-seq (Figure 1H), the region containing the resident, fluorescently-labelled cells in the mature organizer was first excised whole from the embryo, which was then subdivided into the brightly fluorescent regions of the anterior primitive streak and the faintly- or non-labelled regions. Operated embryos were kept at room temperature for 2-4 hours, conditions under which embryos are in developmental diapause but healing can take place, and then placed in a humidified box at 38ºC and allowed to grow.

### Gene expression analyses

RT-PCR (Figure 1J) was done using Phusion High-Fidelity (ThermoFisher, cat. # F553L) a cDNA generated from 5 nodes dissected at stages HH 4-7. The list of primers used The PCR fragments were gel-purified and sequenced to check their sequence corresponds to the intended transcript, and DIG-RNA probes were synthesised using the SP6 promoter included in the reverse primers, as described in <u>GEISHA</u>. In situ hybridizations were performed using the methods described earlier^24^.

### RNA-seq

Total RNA was column-extracted from freshly dissected or laser-captured (LCM) material using RNeasy Micro Kit (Qiagen, cat. # 74004), with a yield of 1-8 ng per sample. Several fixation methods were compared, of which we found Methacarn (methanol:chloroform:acetic acid, 6:3:1 volume proportions) to be most suitable in terms of morphology preservation and quality of extracted RNA. Embryos with the mature Organizer fluorescently labelled were grown in vitro (see above). At the end of incubation, the glass ring with vitelline membrane supporting the embryo was lifted from the culture dish, the membrane and embryo quickly washed with Pannett-Compton solution, and placed in a watch glass over a small pool of ice-cold fixative; fixative was also used to flood the inside of the glass ring and submerge the embryo. The assembly was kept on ice for 30 minutes. Dissected embryos were brought into absolute ethanol, embedded in paraffin and serially cut at 5 µm. Sections were placed on metal framed, 0.9 µm thick POL-membranes (Leica, REF 11505191), and the regions of interest were dissected using a Leica LMD 600 laser capture microdissection system fitted with a x63 lens and fluorescence optics. The total RNA was quality-checked on a pico chip (Bioanalyzer 2100, Agilent), amplified with SMARTer kit (Clontech), and sequenced on an Illumina HiSeq 4000 platform (100bp runs, pair end).

Gene counts and comparisons of gene expression levels between each pair of samples were performed using the Trapnell pipeline^25^. For each pair, we assigned a score of 1 to genes at least 5 times more expressed in the “red” sample (from the MSZ) than in the “white” one (from surrounding regions), and a score of 2 if there was a significant gene count in the “red” sample but zero in the “white” one. Genes were then ranked based on the aggregate score (sheet 1 in Supplementary Table 1). For further assessment by RT-PCR (Supplementary Table S1, sheet 2), we discarded the genes whose expression patterns were known not to be specific to the regions of interest, but included low-ranking genes showing very dissimilar levels in at least some pairs of samples.

## Author Contributions

OV designed and directed the research. TRW, IK and OV performed the experiments. OV wrote the manuscript.

## Acknowledgments

We thank the High-Throughput Genomics Group at the Wellcome Trust Centre for Human Genetics (funded by Wellcome Trust grant reference 090532/Z/09/Z and MRC Hub grant G0900747 91070) for the generation of the Sequencing data. We are grateful to Alfonso Martinez-Arias, Claudio D. Stern, Ben Steventon and Anestis Tsakiridis for comments on the early draft of the manuscript. This word could not have been completed without help from C Linker.

## Figure Legends

**Supplementary Video 1**. 13h time-lapse movie of an embryo with the anterior and middle primitive streak fluorescently labelled (red) from the end of gastrulation (HH stage 3^+^).

**Supplementary Table 1**. Analysis of deep sequencing results.

**Supplementary Table 2**. Primer sequences used for RT-PCR and in situ hybridization.

